# Spectral characterisation of short-wave infrared (SWIR) tissue chromophores and tissue-mimicking phantom optical properties

**DOI:** 10.64898/2026.07.07.736740

**Authors:** Melissa J. Watt, Layla Malouf, Ran Tao, Isabelle Racicot, Thomas R. Else, Janek Gröhl, Sarah E. Bohndiek

**Affiliations:** University of Cambridge, Department of Physics, JJ Thomson Avenue, Cambridge, United Kingdom; Cancer Research UK Cambridge Institute, Robinson Way, Cambridge, United Kingdom; Institute for Experimental Molecular Imaging, University Hospital RWTH Aachen, Aachen, Germany

**Keywords:** short-wave infrared, tissue-mimicking phantoms, optical modelling, haemoglobin, optical properties, integrating sphere

## Abstract

**Significance:** Short-wave infrared (SWIR) sensors promise to expand the capabilities of optical sensing technologies, but the lack of robust data characterising tissue-constituent optical properties in the SWIR makes instrument design challenging.

**Aim:** We characterise and evaluate the optical properties of the dominant chromophores in tissue and tissue-mimicking phantoms, from visible to SWIR wavelengths.

**Approach:** Using single-integrating sphere systems, we measured the optical properties of single-component chromophores (H_2_O, haemoglobin, corn oil, synthetic melanin) and multi-component tissues (whole blood, lard), to decouple contributions from optical scattering, H_2_O absorption and other contributing chromophores; we also characterised commonly-used phantom materials and investigated their potential to mimic soft tissues in the SWIR range using simulations.

**Results:** We provide a consistent dataset of absorption and reduced scattering coefficients (*µ_a_* and *µ_s_*′) that characterise the dominant tissue chromophores from 450 nm out to 1600 nm. These results were shown to be consistent with literature data, where available. We integrate these data into an open-source Python toolkit, SIMPA, for optical modelling and demonstrate soft tissue simulations that can be probed continuously from visible to SWIR wavelengths. Our findings are compared with tissue-mimicking phantoms, highlighting a need for additives for polymer-based phantoms that mimic SWIR water absorption.

**Conclusions:** By providing this open-source dataset, we aim to enable future studies exploring SWIR light-tissue interactions that facilitate rapid assessment and prototyping of next-generation spectroscopy and imaging techniques.

## 1 Introduction

Optical technologies have become ubiquitous tools in healthcare, exploiting the interactions of light with tissue to reveal disease-specific contrast.^1^ Many technologies rely on the absorption of visible (400 – 700 nm) and near-infrared (NIR, 700 – 1000 nm) light by haemoglobin to distinguish disease.^2–4^ For example, in endoscopy, contrast for early cancer arises from the distortion of the vascular architecture, which changes the total haemoglobin absorption in the visible range.^5, 6^ In pulse oximetry, abnormal blood oxygen saturation can be quantified based on the differential optical absorption of oxygenated and deoxygenated haemoglobin, often using visible and NIR wavelengths.^3, 7, 8^ The ubiquity of visible and NIR light in diagnostics and therapeutics has been enabled by the drive towards low-cost, high-quality light sources and detectors in the consumer electronics and photonics industry, accelerating the production of robust and affordable medical devices.^9, 10^

Conversely, the short-wave infrared (SWIR) wavelength range (1000 – 2000 nm) has been relatively underexplored, due to the historically high cost and poor performance of SWIR technologies.^11^ In recent years, however, promising new avenues to realising SWIR medical devices have emerged, including next-generation InGaAs detectors^12, 13^ and upconversion methods that can convert SWIR to visible light.^14–16^ SWIR light emitting diodes (LEDs) are also coming of age, with performance reaching towards those in the visible and NIR.^17, 18^ The use of SWIR light in biomedicine has several potential advantages. First, optical scattering in tissue diminishes rapidly from the NIR to SWIR wavelength range, opening up the possibility of probing deeper into tissue with enhanced spatial resolution.^19, 20^ Second, the SWIR range includes the second and third optical windows of lower water absorption.^21^ Third, aside from the aforementioned optical windows, the dominant tissue chromophores in the SWIR range are water and lipids, opening new biomarker possibilities.^22–24^ Finally, melanin absorption diminishes in the SWIR range, which may be valuable to overcome skin tone bias challenges,^25, 26^ such as those highlighted during the COVID-19 pandemic.^27^

The availability of new sensors and light sources that could underpin SWIR medical devices motivates a fresh look at the potential of this wavelength range in medical imaging,^11^ however, the available data regarding tissue optical properties in the SWIR range are limited.^25, 28^ The lack of robust available data measured on common instruments with consistent processing prohibits the simulation studies needed to define new capabilities provided by the SWIR range and to design prototype instruments.^29, 30^ Here, we undertook a detailed analysis of the SWIR absorption and scattering properties of dominant tissue constituents: haemoglobin, lipids, melanin and water. We measured and report the optical properties of both single-component chromophores (H_2_O, haemoglobin, corn oil, synthetic melanin) and mixtures (whole blood, lard), to decouple contributions from optical scattering, H_2_O absorption and other chromophores with overlapping absorption properties. We then characterised commonly used phantom materials to establish their tissue-mimicking potential in the SWIR range. Through comparison with literature, we showed that our results provide a consistent dataset of absorption (*µ_a_*) and reduced scattering (*µ_s_*′) spectra, characterising the dominant tissue chromophores from 450 nm to 1600 nm. We collated these data for integration with the open-source Python toolkit, SIMPA, for optical modelling^29^ and demonstrated that simulated tissues can be probed continuously from visible to SWIR wavelengths using our dataset, comparing our findings with tissue-mimicking phantoms. By providing this open-source dataset, we aim to enable future studies exploring SWIR light-tissue interactions for the development of the next generation of optical healthcare technologies.

## 2 Materials and Methods

### 2.1 Optical characterisation with single integrating sphere (SIS) systems

We sought to establish the optical properties of the dominant chromophores in tissue (haemoglobin, melanin, water and lipids) and associated tissue-mimicking materials spanning the visible, NIR and SWIR wavelength range (Fig. 1A). The main optical properties of tissue are the absorption coefficient (*µ_a_*), scattering coefficient (*µ_s_*), anisotropy factor (*g*) and reduced scattering coefficient (*µ_s_*′)^31^ given by

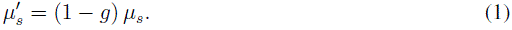

**Fig 1.**
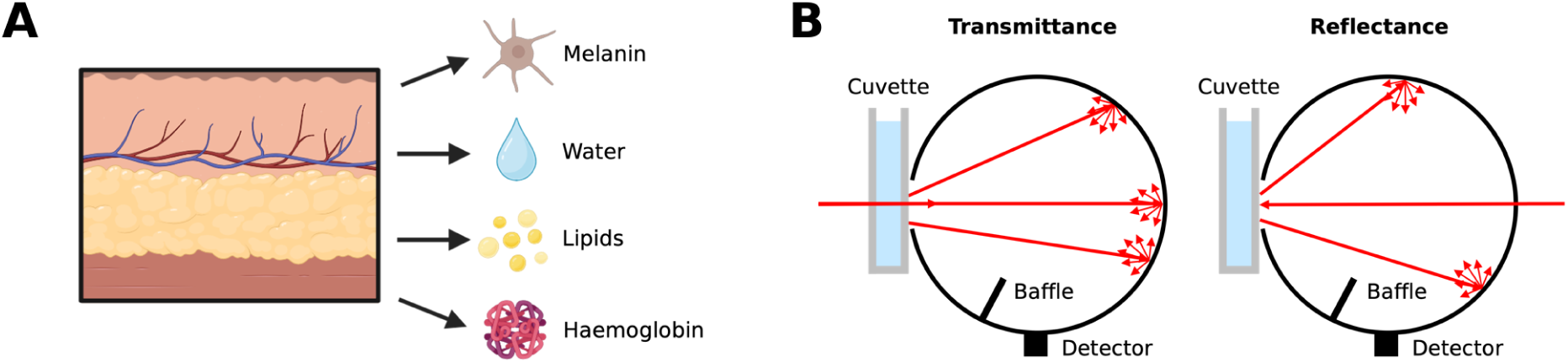
Absorption and scattering coefficients (*µ_a_* and *µ_s_*) of dominant tissue chromophores can be estimated from visible to short-wave infrared wavelengths from measurements of transmittance and reflectance using single integrating sphere (SIS) systems. A) Dominant chromophores in tissue. Created in BioRender. Bohndiek, S. (2026) https://BioRender.com/urkzunj B) Schematic of a single integrating sphere and cuvettes showing transmittance and reflectance. Adapted from Tao *et al.*^38^

Optical absorption and scattering coefficients were obtained using measurements of reflectance and transmittance from single integrating sphere systems (Fig. 1B).

For the case of non-scattering materials, *µ_a_* was estimated by measuring the transmittance of light through sample-containing cuvettes and then applying the Beer-Lambert law

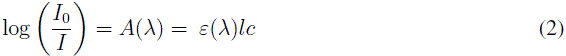

where the ratio of the initial intensity *I*_0_ and measured intensity *I* is equal to the absorbance *A*(*λ*), which is proportional to the path length *l* and to the concentration *c* of the absorber by the molar extinction coefficient *ε*(*λ*).^32, 33^ To cancel the effects of optical interactions due to refractive index (*n*) mismatch between the sample, cuvette glass, and air, measurements were taken through two sample-containing cuvettes of different path lengths *l*_1_ and *l*_2_.^34, 35^ Since

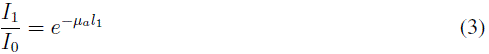

and

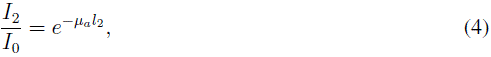

where *I*_1_ and *I*_2_ are the transmittances through cuvettes 1 and 2,^36^ respectively, Eqs. (3) and (4) and can then be rearranged for

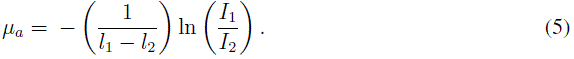

Employing the Beer-Lambert law for *µ_a_* estimation relies on the assumption that the sample does not experience bulk scattering, limiting this method to the characterisation of non-scattering me-dia. A UV-VIS-NIR Spectrophotometer (Shimadzu UV-3600i Plus) with Integrating Sphere Attachment (Shimadzu ISR-603) was used to measure transmittance for estimations of *µ_a_* in non-scattering samples from 250 – 2500 nm. Transmittance measurements were recorded at 2.0 nm intervals using the following settings: High speed scan speed, Transmittance spectrum type, 20 nm slit width, light source switch wavelength of 300 nm, PM-InGaAs Detector Switch Wavelength of 900 nm, InGaAs-PbS Detector Switch Wavelength of 1650 nm and Grating Switch Wavelength of 950 nm with Stair Correction. Measurements on this system are referred to as ‘Shimadzu SIS’ in the remainder of the paper.

For scattering samples, the inverse adding doubling (IAD) algorithm^37^ was employed to calculate *µ_a_* and *µ_s_*′ from integrating sphere reflectance and transmittance measurements, assuming constant *n* and *g* values sourced from literature. The IAD algorithm iteratively optimises an initial guess for *µ_a_* and *µ_s_*′, solving the radiative transport equation to calculate the corresponding total reflectance and transmittance, accounting for integrating sphere specifications. The iteration continues until the calculated values of reflectance and transmittance match the experimentally-measured values.^37, 38^ The Shimadzu system was only able to measure 10 mm wide cuvettes, allowing the loss of scattered photons at the cuvette edges, which would reduce the accuracy of the IAD method, and so for all scattering samples, a custom SIS system developed by Ran Tao was used to estimate *µ_a_* and *µ_s_*′. Samples were positioned flush to the SIS port and illuminated with three halogen light sources (Avantes AVALIGHT-HAL-S-MINI) connected to the sphere with multimode fibre optic patch cables (Thorlabs, M37L01). Transmittance and reflectance measurements were recorded from 450 – 1600 nm, with the visible-NIR spectrometer (Ocean Optics, QEPRO) sampling every 0.7 – 0.8 nm and the SWIR spectrometer (Ocean Optics NIRQ512) recording every 1.6 nm. After the IAD algorithm, the spectra from the visible-NIR and SWIR spectrometers were stitched together by averaging in their overlapping region (950 – 1000 nm); the stitched spectra were then resampled to 2 nm intervals. Measurements on this system are referred to as ‘in-house SIS’ for the remainder of the paper.

### 2.2 Preparation of chromophore samples

Chromophore samples measured with the in-house SIS system were dispensed into custom fused silica cuvettes (FireflySci, Suprasil 3001 silica, 1.5 mm pathlength, 1.25 mm wall thickness, 45*×*45 mm area, refractive index of 1.4545), which housed 2.7 mL aliquots. For samples measured in the Shimadzu SIS, 1 mm path length (Hellma UK Ltd 100-1-40, 100-QS, PL 1 mm) and 2 mm path length (Hellma UK Ltd 100-2-40, 100-QS, PL 2 mm), 10 mm wide cuvettes, loaded with 350 µL and 700 µL of sample, respectively, were used. Cuvettes were cleaned with a 1 % Hellmanex III (Sigma Aldrich, Z805939-1EA) in deionised (DI) water (H_2_O) solution, followed by ethanol, then acetone, and dried with nitrogen gas, between samples. Unless otherwise specified, all data collection consisted of three technical replicates of three independently-prepared samples.

#### 2.2.1 Solvents

Samples of DI H_2_O, deuterium oxide (D_2_O, Sigma Aldrich, CAS: 7789-20-0) and dimethyl sulfoxide (DMSO, Biomol GmbH, CAS: 67-68-5) were prepared for characterisation with the Shimadzu SIS. The D_2_O and DMSO were dispensed directly into the 1 mm and 2 mm Hellma cuvettes, whilst the stock of DI H_2_O (sourced from the central building supply) was filtered with a 0.22 µm syringe filter to remove any impurities before being loaded into cuvettes. Being standard and stable solvents, the three independent replicates of H_2_O and DMSO were measured just once.

#### 2.2.2 Haemoglobin

We extracted haemoglobin from whole horse blood in ethylenediaminetetraacetic acid (EDTA, TCS Biosciences HB073). The haematocrit of the whole blood was estimated by comparing the plasma-pellet volume ratio after spinning three 5 mL samples of whole blood in 15 mL centrifuge tubes (Sarstedt) in a cooled centrifuge at 4 ^◦^C for 10 minutes at 500 rcf. The volume of the red blood cell (RBC) pellets in the three tubes were 2.5 mL, 2.0 mL and 2.7 mL ± 0.1 mL, giving an average of 2.4 ± 0.4 mL, or an estimated haematocrit of 48 ± 8%, which is consistent with the typical 40 – 45 % expected for human adults.^39–41^

Haemoglobin samples were prepared by adapting the protocol for cell lysis and extraction of haemoglobin from whole human blood by Gruensfelder *et al.*^28^ We deviated from this protocol in two notable stages: in the first, we washed the blood samples in an isotonic buffer (9% NaCl (Fisher Scientific, CAS: 7647-14-5) in D_2_O), which prevents red blood cell lysis when removing plasma and platelets, improving haemoglobin yield. In the second, we used Size Exclusion Chromatography (SEC) to isolate haemoglobin from the cell lysate, rather than syringe filtration. This ensures that only the desired haemoglobin is retained, rather than all lysate products below the mesh size of the syringe filter. We used 0.4 g Sephadex G-50 (Sigma Aldrich, CAS: 9048-71-0) as the size exclusion resin, swollen in 12 mL D_2_O at 50^◦^C for approximately 100 min. The resin slurry was poured into a 5 mL polypropylene column (Thermo Scientific, PN: 29922) and left to settle for approximately 1 hour before using for SEC.

To prepare the haemoglobin solution for SEC, the plasma was removed from the centrifuged samples used for haematocrit estimation, and then the RBC pellet was diluted back to 5 mL with the isotonic buffer. The cells were washed twice more, as per the protocol, before the buffer was removed one final time. Then, 1 mL of the RBC pellet from each tube was drawn and dispensed into three fresh tubes. The RBCs were lysed by introducing 4.5 mL of D_2_O into each tube and inverting several times to mix. Two samples of 200 µL of the resulting haemoglobin solution were then filtered through the SEC column for each of the three tubes (the column was flushed with D_2_O between each sample). The most concentrated fractions were then combined, yielding three sets of haemoglobin solution.

To prepare oxygenated samples for optical characterisation, 800 µL of haemoglobin solution was combined with 800 µL of D_2_O and inverted 10 times to mix. To prepare deoxygenated samples, a stock solution of 35 mg sodium dithionite (Sigma Aldrich, CAS: 7775-14-6) in 6 mL D_2_O was prepared for chemical deoxygenation. Then, 700 µL of haemoglobin solution was then combined with 700 µL of the dithionite solution and inverted 10 times to mix. Immediately after mixing, the oxygenated and deoxygenated haemoglobin solutions were flushed through an optical oxygen meter (Pyroscience FireSting-O2) to measure the partial pressure of oxygen (pO_2_), before being loaded into three sets of 1 mm and 2 mm path length cuvettes and immediately measured with the Shimadzu SIS. To determine the sO_2_ of the solutions, the minimum and maximum pO_2_ values (measured in mmHg) of the deoxygenated and oxygenated samples, respectively, were calculated. The Severinghaus equation^42^ was used to convert to sO_2_, yielding sO_2_ = 99% for the oxygenated haemoglobin solutions and sO_2_ = 0% for the deoxygenated solutions.

#### 2.2.3 Whole Blood

We characterised whole horse blood in EDTA (TCS Biosciences HB073) to enable the quantification of whole blood scattering and to evaluate the contributions from water in whole blood absorption. To avoid sensor saturation arising from strong absorption by haemoglobin at visible wavelengths, oxygenated an deoxygenated samples of both whole horse blood and 1 % solutions of the whole horse blood in phosphate-buffered saline (PBS, Gibco, PN: 10010-015) were prepared for optical characterisation. Characterisations were conducted with the same batch of blood as described in Sec. 2.2.2, apart from for whole (undiluted) oxygenated blood where characterisations were conducted with a new batch of whole horse blood in EDTA, assumed to be almost fully oxygenated (sO_2_ *>* 98 %), to be consistent with sample age. Since we may expect slight batch-to-batch variation in haematocrit of our whole horse blood samples, we again estimated haematocrit as described previously. These data show an estimated haematocrit of 47 ± 8%, consistent both with the previously-used batch of whole horse blood, and with the typical 40 – 45 % expected for human adults.^39–41^

For oxygenated blood, three cuvettes were filled with aliquots of the whole blood, drawn from the bottom of the container after inverting, before being taken for measurement on the in-house SIS. For the 1 % oxygenated solution, three independently-prepared replicates of 0.1 mL whole horse blood were added to 9.9 mL PBS. An sO_2_ of 99 % was measured with an oxygen meter (Pyroscience FireSting-O2, see Sec. 2.2.2) before the solution was loaded into cuvettes for measurement with the in-house SIS. Deoxygenated samples were prepared using sodium dithionite to chemically deoxygenate three samples of whole blood (25 mg sodium dithionite in 10 mL blood, inverted several times to mix). An sO_2_ of 0 % was measured; for reference, three replicates of 10 mL of whole blood (not deoxygenated) were also flowed through the oxygen meter giving sO_2_ = 98 %. The blood was quickly loaded into the cuvettes and immediately measured using the in-house SIS. For the 1 % deoxygenated solution, three independently-prepared replicates of 0.1 mL whole horse blood were added to 9.9 mL PBS, deoxygenated with 25 mg sodium dithionite and inverted to mix. The pO_2_ of the three samples was measured to be 0 % before being loaded into the cuvettes for immediate characterisation. Consistency of oxygenation was verified for all samples.

For the undiluted blood samples, *g* was assumed to be 0.98 and *n* assumed to be 1.35 for the IAD algorithm, whilst for the 1 % solutions, *g* was assumed to be 0.9998 and *n* assumed to be 1.33. After *µ_a_* estimation, *µ_a_* of the 1 % solutions from 450 – 600 nm was multiplied by a factor of 100 (to account for the 100*×* dilution) and stitched to the *µ_a_* estimations of the whole (undiluted) blood from 600 – 1600 nm.

Flow cytometry was used to measure the horse RBC shape and size for comparison with human RBCs. A fresh sample of whole horse blood in EDTA was diluted 1000*×* with PBS (Fisher Bioreagents, 10X Phosphate Buffered Saline BP399-20, diluted to 1 *×* and autoclaved) and measured with an imaging flow cytometer (Cytek Amnis ImageStream*^X^* Mk II). The Cytek IDEAS Image Analysis Software was used to analyse the brightfield RBC images: the in-built cell masking and area calculation functions were implemented and cell diameters were determined as twice the square root of the calculated area divided by pi. The RBC images were gated into four population types: first, aspect ratio against areas was used to sort side view from face view cells, then rounded cells were separated from star-shaped cells by examining a map of Haralick^43^ (H) variance standard deviation against H variance mean, and finally, cells with a flat surface were distinguished from those with a concave centre using a map of H homogeneity mean against maximum raw pixel values. Additionally, a haematology analyser (Cormay Diagnostics Mythic 5Vet PRO) was used to measure the haematocrit of the batch of horse blood examined with flow cytometry. Three measurements were taken, yielding 44.5 %, 44.2 % and 43.8 %, consistent with estimated haematocrit obtained from RBC pellet and plasma volumes, as described previously.

#### 2.2.4 Lipids

To compare the absorption features of animal and plant-derived lipids, samples of lard (Tesco Ireland Ltd.) and corn oil (Sigma Aldrich, C8267-500ML) were prepared. Lard (reduced pig fat) was heated with a microwave until melted and then loaded into cuvettes for characterisation with the in-house SIS. The lard samples were left at room temperature to set, remelted in a water bath where necessary to release trapped air bubbles, and were measured once solid. For lard, *g* was assumed to be 0.7 and *n* was assumed to be 1.47 for the IAD algorithm. As corn oil is non-scattering, the Shimadzu SIS was used for characterisation.

#### 2.2.5 Synthetic Melanin

Solutions of synthetic melanin (Sigma Aldrich, CAS: 8049-97-6) in DMSO (Biomol GmbH, CAS: 67-68-5)) with a concentration of 0.21 mg/mL were prepared. Three stock solutions composed of 21 ± 1 mg of synthetic melanin in 10 mL DMSO were prepared and then diluted further by a factor of 10. Each solution was inverted several times and vortex mixed for 10 seconds to ensure that melanin was dissolved before being taken for measurement with the Shimadzu SIS.

#### 2.2.6 Data Processing and Statistics

All data processing was performed in Python. First, *µ_a_* and *µ_s_*′ (for scattering samples) were estimated for each replicate, as detailed in Sec. 2.1. Then, separately for each chromophore, the *µ_a_* spectra were pooled so that the mean and standard deviation spectra could be calculated, and similarly for *µ_s_*′. To remove detector and noise artefacts, infinities and values of *µ_a_* less than 0.001 cm^−1^ were removed from all spectra. Values where *µ_a_* ± three standard deviations of *µ_a_* was less than 0.001 cm^−1^ (the lower limit of detection of the Shimadzu SIS) were removed to account for artefacts due to low sensitivity. To enable quantitative evaluation of our measured optical properties, we compared our results against references from literature using spectral angle mapping (SAM), Pearson’s correlation coefficient (R) and root mean square error (RMSE). The measured and reference spectra were interpolated to match the step size of the spectrum with the largest wavelength step size for each chromophore. The SAM score, *θ* is defined as

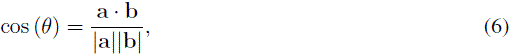

where **a** and **b** are the measured and reference spectra, respectively. R, and associated 95 % confidence interval, were determined using the stats. pearsonr function in the SciPy^44^ library. The RMSE was calculated using the NumPy^45^ library as

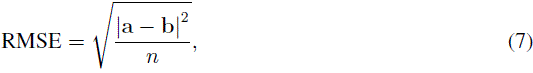

where **a** and **b** are the measured and reference spectra, respectively, and *n* is the number of spectral points. We estimate the uncertainty for each statistic, by computing half the distance between the lower and upper bound. We calculate these bounds by computing the measures from the mean spectrum ± the wavelength-dependent standard deviation. Statistical analyses were performed between 450 – 1000 nm and 1000 – 1600 nm to allow visible-NIR to SWIR comparisons and to enable inter-chromophore comparison.

### 2.3 Tissue-Mimicking Phantoms

Tissue-mimicking phantoms were fabricated for comparison with the biomolecule optical properties. Recipes describing protocols for fabricating agar,^46^ polydimethylsiloxane (PDMS)^47^ and copolymer-in-oil^48^ phantoms were selected from highly-cited papers and used to fabricate reference phantoms. Custom 3D-printed tough polylactic acid (PLA) moulds were designed to create thin rectangular slabs of the agar and copolymer-in-oil phantom materials so that the optical absorption and scattering properties of each material could be characterised using the in-house SIS system. A mould that forms 60 *×* 40 mm wide phantoms with a thickness of 2 mm was used for the co-polymer-in-oil phantoms. As the agar phantoms are hydrogels, square (40 mm outer sides, 38 mm inner sides, 1.5 mm deep) tough PLA frames containing the phantom were slotted into a custom sample holder with quartz windows to prevent contamination of the SIS system. The PDMS phantoms were poured into 90 mm plastic petri dishes for compatibility with the vacuum chamber. Three slabs of each material were poured and characterised, with spectral processing as described in Sec. 2.2.6.

The agar phantoms contained India ink (Rotring Drawing Ink) and Intralipid (Sigma Aldrich, CAS: 68890-65-3) for optical absorption and scattering. First, 2.0 g agar (Sigma Aldrich, CAS: 9002-18-0) was mixed with 198 mL DI water and heated with a microwave on the high setting for approximately 100 seconds until boiling. Then, the mixture was stirred continuously and monitored with a digital thermometer. Once cooled to 60 ^◦^C, 2.0 g of Intralipid was stirred into the agar solution along with 2 µL of India ink. When the phantom material reached 40 ^◦^C, approximately 2.2 mL of material was dispensed into each of the three PLA frames using a pipette and left to set at room temperature for 5 minutes before characterisation with the in-house SIS. The thickness of the phantoms was assumed to be 1.5 mm, whilst a *g* of 0.9 and an *n* of 1.33 were assumed for the IAD algorithm.

The PDMS phantoms used India ink (Rotring Drawing Ink) and titanium dioxide (Sigma Aldrich, CAS: 1317-70-0) to provide scattering and absorption. A silicone elastomer (Dow, Sylgard 184 Silicone Elastomer) was used as the PDMS matrix – this is a two-part elastomer, comprising a base material (Part A) and the curing agent (Part B). Titanium dioxide (40 mg) was dispersed in 5 mL of curing agent by sonicating in a water bath for 30 minutes, at room temperature. The India ink (14 µL) was mixed into the PDMS base (48.7 g) by stirring manually for 10 minutes. Then, the 5 mL solution of titanium dioxide in curing agent was added to the pigmented PDMS base, and the mixture was again stirred manually for 15 minutes to combine. The mixture was then poured into three 90 mm petri dishes, to a depth of approximately 2 mm, before being placed in a vacuum chamber for 20 minutes to remove entrapped air bubbles. The phantoms were cured at room temperature for 2 days before characterisation. Their thicknesses was measured for the IAD algorithm using digital callipers to be 2.24 ± 0.01 mm, 2.18 ± 0.04 mm and 2.35 ± 0.04 mm; *g* was assumed to be 0.9 and *n* was assumed to be 1.40.

For the copolymer-in-oil phantoms, 1000 µL of a 10 mg/mL stock solution of alcohol-soluble nigrosin (Sigma Aldrich, CAS: 11099-03-9) in mineral oil (Sigma Aldrich, CAS: 8042-47-5) alongside 0.15 g of titanium dioxide (Sigma Aldrich, CAS: 1317-70-0) were mixed with 83.8 g of mineral oil and sonicated in a water bath for 60 minutes at 90^◦^C. Then, the mixture was combined with 25.3 g poured 25.3 g of polystyrene-block-poly(ethylene-ran-butylene)-block-polystyrene (Sigma Aldrich, CAS: 66070-58-4) and 1.68 g of low density polyethylene (Alfa Aesar, CAS: 9002-88-4).

The mixture was heated to 160 ^◦^C in a silicone oil bath under continuous magnetic stirring until melted. The material was degassed in a vacuum chamber and poured into the 3D printed moulds. Once set, the phantom thicknesses were measured using digital callipers to be 2.30 ± 0.03 mm, 2.51 ± 0.01 mm and 2.64 ± 0.10 mm; *g* was assumed to be 0.7 and *n* was assumed to be 1.43 for the IAD algorithm.

### 2.4 Computational Models

We collated our measured tissue chromophore spectra into a single continuous dataset of optical properties from visible to SWIR to use them for tissue simulations. Computational tissue models of muscle and fat were constructed using the open-source *simulation and image processing for photonics and acoustics* (SIMPA) Python toolkit.^29^ SIMPA models photon propagation using the Monte Carlo method in optical imaging simulations by interfacing with the Monte Carlo eXtreme software package.^49^ SIMPA houses libraries of biological structures, morphological geometries, and tissue optical properties for simulations between 450 – 1000 nm, which were built upon to incorporate the SWIR properties collected in this work.

In order to simulate soft tissues, including blood, muscle and fat, out to SWIR wavelengths, we replaced SIMPA’s absorption data for oxygenated and deoxygenated blood and fat with our measurements of horse blood and lard. Due to the lower limit of detection of our instrument, we were unable to determine the spectral properties of lard accurately in the visible and NIR range, so we stitched the absorption coefficient of pig fat by van Veen *et al.*^50^ from 450 – 1000 nm to our lard *µ_a_* from 1000 – 1600 nm. Additionally, the absorption coefficient of water by Segelstein^23^ was used as this dataset already characterises water continuously over the visible and SWIR range. For scattering, we implemented models for blood, muscle, and fat scattering using Rayleigh and Mie theory:

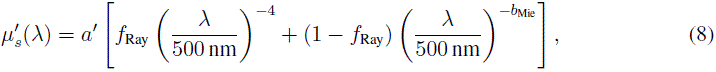

where *a*^′^ = *µ_s_*′,_500*nm*_, the empirically measured value of *µ_s_*′ for a tissue at 500 nm, *f*_Ray_ is the fraction of Rayleigh scattering and *b*_Mie_ is the Mie scattering power (Table 1).^36^

**Table 1.** Coefficients for modelling reduced scattering coefficient of tissues using Eq. (8). ^∗^*a*^′^ for blood and muscle converted from *µ_s_* to *µ_s_*′ using Eq. (1) assuming *g* = 0.98 and *g* = 0.9 for blood and muscle, respectively.

| Tissue | $a' = \mu'_{s,500 \text{ nm}} [\text{cm}^{-1}]$ | $f_{\text{Ray}} [\text{a.u.}]$ | $b_{\text{Mie}} [\text{a.u.}]$ | Reference |
| --- | --- | --- | --- | --- |
| Blood | 23.4* | 0.0 | 0.93 | <sup>29</sup> |
| Muscle | 16.1* | 0.21 | 1.50 | <sup>29,51</sup> |
| Fatty tissue | 19.3 | 0.174 | 0.447 | <sup>29,36</sup> |

It should be noted that the blood scattering coefficients are those optimised by Gröhl *et al.*^29^ where *µ_s,_*_500_ *_nm_* = 1170 cm^−1^ is converted to *µ_s_*′ using Eq. (8), assuming *g* = 0.98. Eq. (8) was also used to model whole human blood scattering for simulation purposes. This was done despite measuring directly whole horse blood to account for the differences in RBC shape between horse and human blood, which leads to slightly different scattering properties (see Sec. 3.2).

The fat model was made using SIMPA’s predefined volume fractions for fat tissue: water (68 % v/v, *µ_a_* by Segelstein^23^ and negligible scattering), blood (1 % v/v, 95 % oxygenation, our horse blood *µ_a_* and blood *µ_s_* as defined in Table 1) and fat (31 % v/v, pig fat *µ_a_* paired with fatty tissue *µ_s_*′, see Table 1). A constant *g* of 0.9 was assumed. The muscle model was also built using SIMPA’s predefined volume fractions for muscle tissue: a custom water molecule (68 % v/v, *µ_a_* by Segelstein^23^ and *µ_s_*′ for muscle as defined in Table 1), blood (1 % v/v, 95 % oxygenation, our horse blood *µ_a_* and blood *µ_s_* as defined in Table 1) and muscle scatterer (31 % v/v, negligible *µ_a_* paired with muscle *µ_s_*′, see Table 1). A constant *g* of 0.9 was also assumed. To bridge the gap between simulation and experiment, we also built models of our tissue-mimicking phantoms by loading phantom optical properties (*µ_a_* and *µ_s_*′ measured with the in-house SIS, assuming *g* = 0.9 for the agar and PDMS and *g* = 0.7 for the co-polymer-in-oil phantoms) into SIMPA as new tissue types (each 100 % v/v). Uniform volumes of 20 *×* 20 *×* 20 mm were defined for each material (muscle, fat, Agar, PDMS and co-polymer-in-oil) in a grid with a voxel size of 100 *µ*m to capture detail on the order of epidermal thickness. The tissue models were illuminated with a pencil beam centred at the surface of the X-Y plane of each volume to propagate down in the Z-direction. Wavelengths ranging from 450 – 1590 nm were probed in 10 nm steps. Per simulation, 5 *×* 10^7^ photons were modelled. All simulations were run on a computer with NVIDIA GeForce RTX 4060, 8GB GDDR6X, HDMI, 3 DP GPU, Intel(R) Core(TM) i9 14th Gen 14900K (36 MB cache, 24 cores, 32 threads, 3.2 GHz to 6.0 GHz, 125W) processor, and 64 GB RAM.

Simulations were processed using the SIMPA toolkit^29^ and the NumPy python package;^45^ additionally, large language models were used to support code development. The effective penetration depth for each model was calculated as where the fluence falls to 37% (1*/e*) of its maximum value. To achieve this, diffusion of the pencil beams through the 3-dimensional volumes were reduced to 1-dimension by calculating the total energy per X-Y plane (by taking the sum of the fluence at each X-Y plane and multiplying by the area of each voxel: 0.01 mm^2^). These values were then normalised by the maximum value of the total energy so that the index corresponding to where the normalised energy is closest to 37% of the maximum value could be determined. The depth in the Z-direction corresponding to the determined index was then found to give the effective penetration depth. This calculation was performed for each wavelength probed.

## 3 Results

### 3.1 Absorption coefficient (µ_a_) estimations show good agreement with reference compounds H_2_O and D_2_O

We first validated the performance of the Shimadzu SIS and in-house SIS systems by comparing our estimation of *µ_a_* of H_2_O using the Shimadzu SIS (solid red) with reference measurements by Segelstein^23^ (dashed grey) and characterisation of H_2_O by Ran Tao using the in-house SIS system (dashed black) (Fig. 2A). Our characterisation compared favourably between 1000 – 1600 nm with Segelstein, yielding a spectral angle of 0.10738 ± 0.00065 rad (Table 2). Similarly, a spectral angle of 0.097 ± 0.003 rad is calculated comparing our measurement with previous measurements by Ran Tao. Tao’s characterisation is also compared with Segelstein yielding a spectral angle of 0.12176 ± 0.00016 rad. We observe strong *R* values *>* 0.99 as well as RMSE values *<* 2 cm^−1^ comparing both Segelsetin and Tao with our measurement using the Shimadzu SIS (Table 2). The analyses of the remaining chromophores characterised in this work are considered relative to these H_2_O statistics due to the consistency of H_2_O spectra in literature^53^ and agreement between our data, measurements by Ran Tao and Segelstein’s H_2_O characterisations. As *R* was consistent across most characterised chromophores and RMSE is susceptible to amplitude differences between measured and reference datasets, we relied on spectral angles to compare our measurements with reference data.

**Fig 2.**
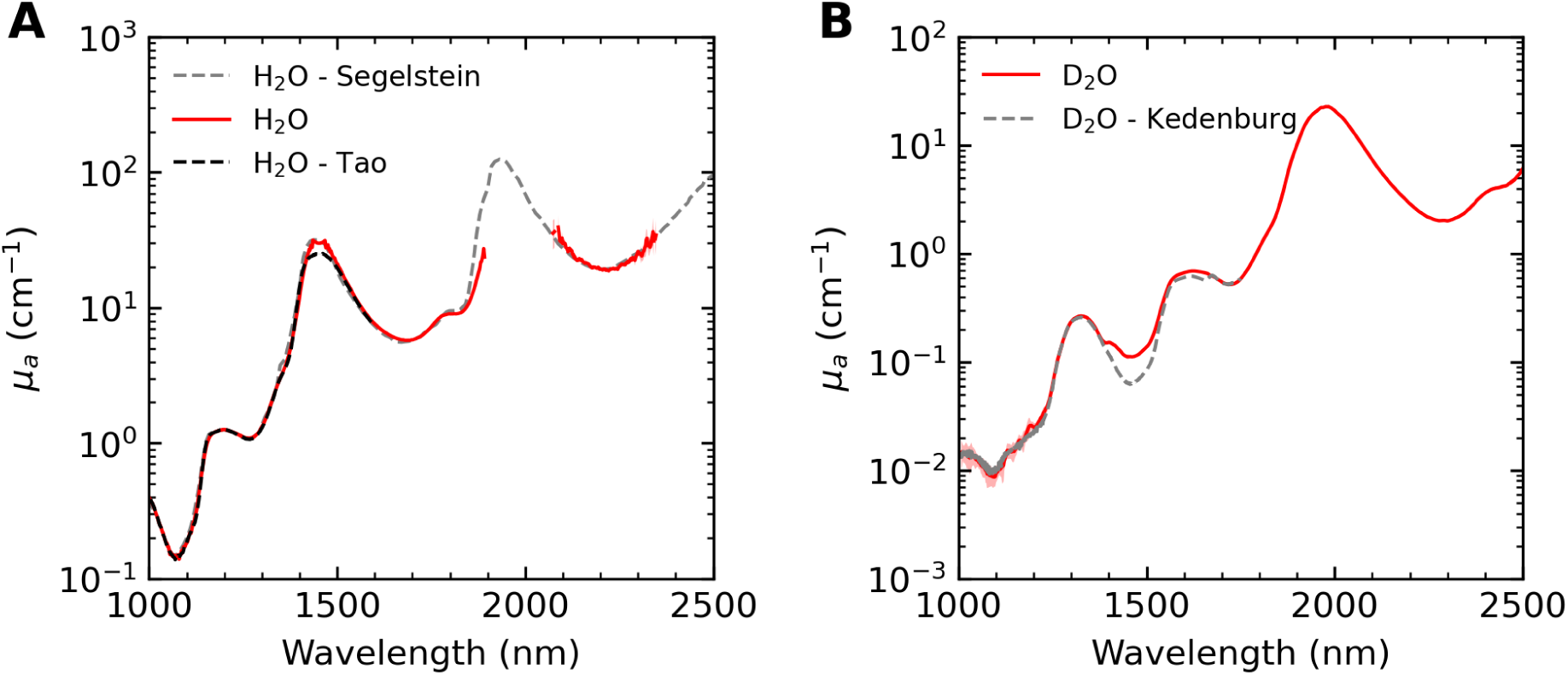
Estimations of the absorption coefficient *µ_a_* of H_2_O and D_2_O are consistent with references from literature from 1000 – 1600 nm. Estimated *µ_a_* of H_2_O (A) and D_2_O (B) calculated using Beer-Lambert analysis of transmittance measurements from the Shimadzu SIS (solid red) are compared to reference data (dashed grey) by Segelstein^23^ (A) and Kedenburg^34^ (B). Estimated *µ_a_* of H_2_O measured by Tao, calculated with reflectance and transmittance measurements from the in-house SIS using the inverse adding doubling algorithm, are presented (dashed black).

**Table 2.**
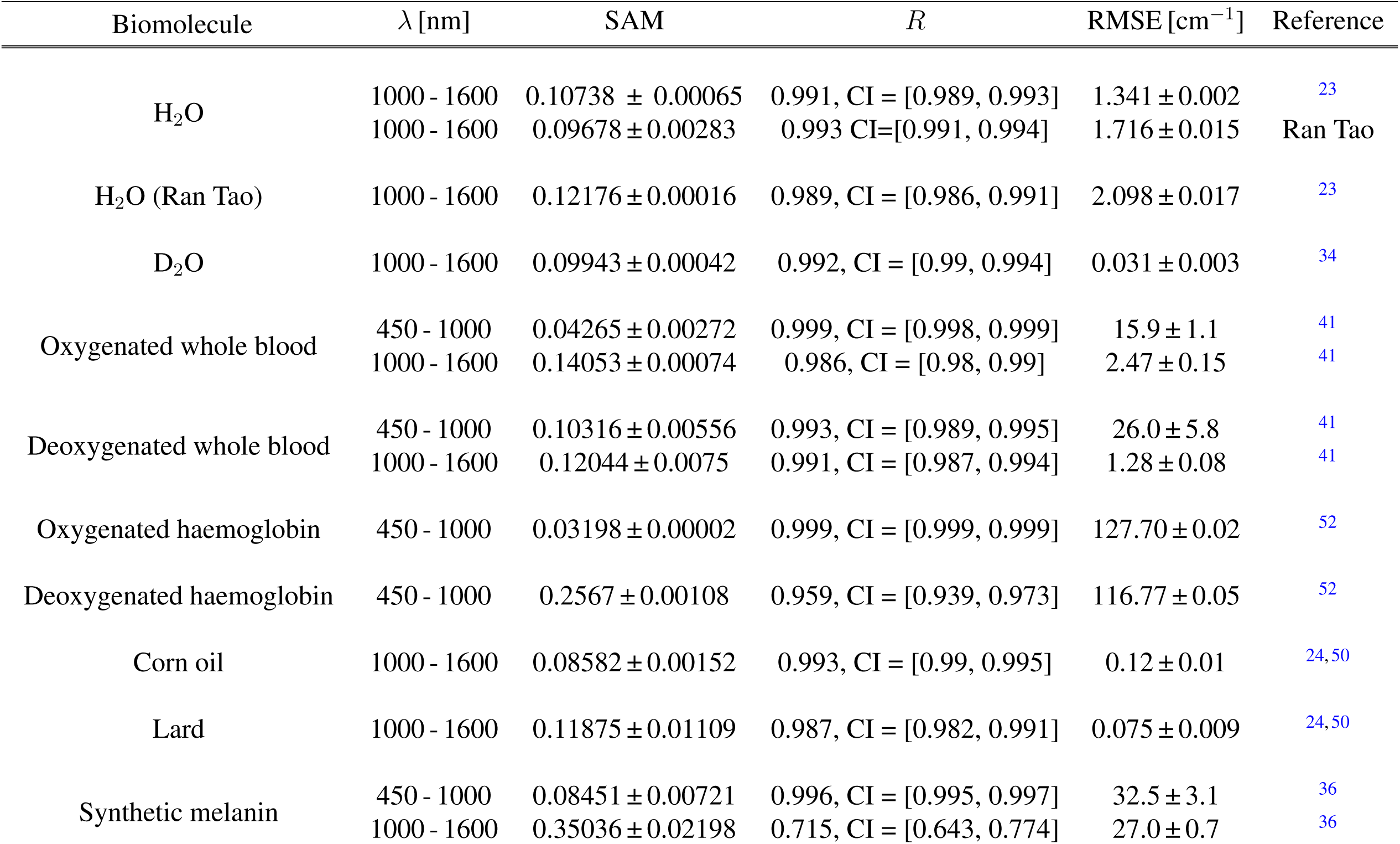
Spectral angle mapping score (SAM), Pearson correlation coefficient (*R*), and root mean square error (RMSE) of measured absorption coefficients compared against references from literature.

| Biomolecule | $\lambda$ [nm] | SAM | $R$ | RMSE [$\text{cm}^{-1}$] | Reference |
| --- | --- | --- | --- | --- | --- |
| H<sub>2</sub>O | 1000 - 1600 | 0.10738 ± 0.00065 | 0.991, CI = [0.989, 0.993] | 1.341 ± 0.002 | <sup>23</sup> |
| | 1000 - 1600 | 0.09678 ± 0.00283 | 0.993 CI=[0.991, 0.994] | 1.716 ± 0.015 | Ran Tao |
| H<sub>2</sub>O (Ran Tao) | 1000 - 1600 | 0.12176 ± 0.00016 | 0.989, CI = [0.986, 0.991] | 2.098 ± 0.017 | <sup>23</sup> |
| D<sub>2</sub>O | 1000 - 1600 | 0.09943 ± 0.00042 | 0.992, CI = [0.99, 0.994] | 0.031 ± 0.003 | <sup>34</sup> |
| Oxygenated whole blood | 450 - 1000 | 0.04265 ± 0.00272 | 0.999, CI = [0.998, 0.999] | 15.9 ± 1.1 | <sup>41</sup> |
| | 1000 - 1600 | 0.14053 ± 0.00074 | 0.986, CI = [0.98, 0.99] | 2.47 ± 0.15 | <sup>41</sup> |
| Deoxygenated whole blood | 450 - 1000 | 0.10316 ± 0.00556 | 0.993, CI = [0.989, 0.995] | 26.0 ± 5.8 | <sup>41</sup> |
| | 1000 - 1600 | 0.12044 ± 0.0075 | 0.991, CI = [0.987, 0.994] | 1.28 ± 0.08 | <sup>41</sup> |
| Oxygenated haemoglobin | 450 - 1000 | 0.03198 ± 0.00002 | 0.999, CI = [0.999, 0.999] | 127.70 ± 0.02 | <sup>52</sup> |
| Deoxygenated haemoglobin | 450 - 1000 | 0.2567 ± 0.00108 | 0.959, CI = [0.939, 0.973] | 116.77 ± 0.05 | <sup>52</sup> |
| Corn oil | 1000 - 1600 | 0.08582 ± 0.00152 | 0.993, CI = [0.99, 0.995] | 0.12 ± 0.01 | <sup>24,50</sup> |
| Lard | 1000 - 1600 | 0.11875 ± 0.01109 | 0.987, CI = [0.982, 0.991] | 0.075 ± 0.009 | <sup>24,50</sup> |
| Synthetic melanin | 450 - 1000 | 0.08451 ± 0.00721 | 0.996, CI = [0.995, 0.997] | 32.5 ± 3.1 | <sup>36</sup> |
| | 1000 - 1600 | 0.35036 ± 0.02198 | 0.715, CI = [0.643, 0.774] | 27.0 ± 0.7 | <sup>36</sup> |

*3.1 Absorption coefficient ( $\mu_a$ ) estimations show good agreement with reference compounds $H_2O$* *and $D_2O$*

Characterisation of D_2_O further supports our system validation: our *µ_a_* estimation using the Shimadzu SIS (solid red) compares favourably with reference measurements by Kedenburg^34^ (dashed grey) (Fig. 2B). Quantitatively, we observe a SAM score of *<* 0.1 when comparing our measurement with Kedenburg between 1000 – 1600 nm in addition to *R >* 0.99 and RMSE of 0.031 ± 0.003 cm^−1^ (Table 2).

### 3.2 Optical properties of whole blood show strong oxygenated haemoglobin absorption below 1200 nm

The optical properties of whole horse blood measured in this study (Fig. 3A) were compared to an average *µ_a_* of oxygenated and deoxygenated human whole blood compiled from literature references by Bosschaart *et al.*^41^ (sO_2_ *>* 98 % and sO_2_ = 0 %, respectively). The oxygenated blood yields a low spectral angle of 0.04265 ± 0.000272 rad when compared with Bosschaart between 450 – 1000 nm (Table 2), which increases to 0.14053 ± 0.00074 rad between 1000 – 1600 nm. The poorer agreement in the SWIR range is likely due to the literature source compiling multiple datasets, which were noted by Bosschaart to be ignored or smoothed in practice.^41^ Conversely, our spectrum is smoother as a result of our characterisation using only one instrument. Analysis of the deoxygenated blood spectrum yields strong SAM scores (0.10316 ± 0.00556 and 0.12044 ± 0.00750 for visible-NIR and SWIR, respectively), which are comparable with the SAM scores of our H_2_O references (Table 2). Again, it is likely that the higher SWIR spectral angle can be attributed to data compilation artefacts in the reference data as above and our measurements more closely following H_2_O absorption after 1000 nm.

**Fig 3.**
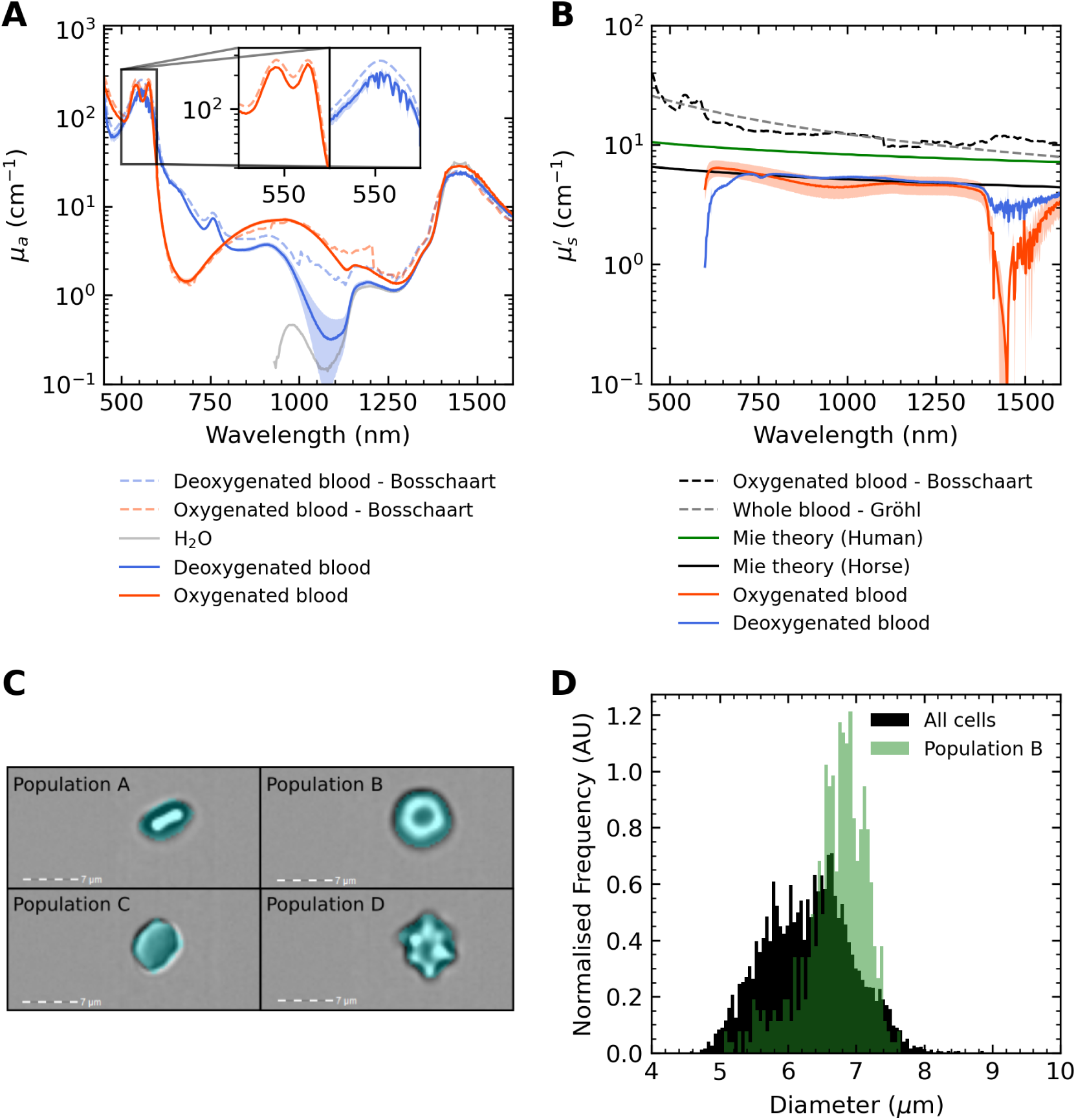
Characterisation of the absorption coefficient (*µ_a_*) of oxygenated and deoxygenated whole horse blood from 450 – 1600 nm and reduced scattering coefficient (*µ_s_*′) from 600 – 1600 nm. A) Estimated *µ_a_* of a 1% solution of whole blood in phosphate-buffered saline (PBS) is presented from 450 – 600 nm, stitched to estimated *µ_a_* of undiluted whole blood from 600 – 1600nm (solid red). Similarly, a 1% solution of deoxygenated whole blood in PBS is presented from 450 – 600nm, stitched to the estimated *µ_a_* of undiluted whole blood from 600 – 1600nm (solid blue). An average *µ_a_* of oxygenated (dashed red) and deoxygenated (dashed blue) human whole blood compiled from literature references by Bosschaart *et al.*^41^ is compared with our estimations for *µ_a_*. B) Estimated *µ_s_*′ of oxygenated and deoxygenated whole horse blood from 600 – 1600nm (solid red and blue, respectively). An average *µ_s_*′ compiled from literature by Bosschaart *et al.*^41^ (dashed black) and an approximation based on Rayleigh and Mie theory using coefficients optimised by Gröhl *et al.*^29^ (dashed grey). C) Brightfield images of horse red blood cells (RBCs) imaged with flow cytometer. Four population types were identified side profile RBCs (A), circular cells with a concave middle (B, assumed to be erythrocytes), flat cells (C, assumed to be spherocytes) and star-shaped cells (D, assumed to be Burr cells). The cyan overlays show the masking function used to measure cell diameter by the flow cytometer software. D) Histogram showing distribution of horse RBC diameters from all RBC populations (black) with population B overlayed (green).

The estimated *µ_s_*′ of oxygenated (sO_2_ = 98%) and deoxygenated (sO_2_ = 0%) whole horse blood from 600 – 1600 nm were compared to an average *µ_s_*′ spectrum calculated using Eq. (1) from *µ_s_* and *g* data compiled from literature by Bosschaart *et al.*^41^ (Fig. 3B). An approximation based on Rayleigh and Mie theory using Eq. (8) with coefficients optimised by Gröhl *et al.*^29^ is also plotted for comparison. Both reference spectra show reduced scattering values arger than those reported in our data, which we attribute to differences in horse and human red blood cell size.

To test this, flow cytometry was used to image and measure the size of RBCs from a sample of horse blood. Representative images of the four identified population types: side profile RBCs (A), circular cells with a concave middle (B, assumed to be erythrocytes), flat cells (C, assumed to be spherocytes) and star-shaped cells (D, assumed to be Burr cells) are presented (Fig. 3C). Analysis of the population distribution of the RBC diameters of all the cell populations (black) yielded a mean diameter of 6.34 ± 0.65 µm (Fig. 3D). For comparison, healthy human RBCs have an average diameter of 7.8 µm,^54^ still higher than the mean diameter of population B, assumed to be the healthy population, which was calculated to be 6.77 ± 0.46 µm. Thus, lower scattering would be expected for horse blood according to Mie theory, due to the smaller RBC diameter (scattering particle) size. We simulated the expected *µ_s_*′ spectra using Mie theory^55^ assuming a human RBC diameter of 7.8 µm (solid green) and a horse RBC diameter of 6.34 µm (solid black) to validate our findings (Fig. 3B), showing lower scattering for horse blood, consistent with our measured *µ_s_*′ spectra.

### 3.3 Water dominates haemoglobin solution absorption in SWIR

Comparing the *µ_a_* of the oxygenated (sO_2_ = 99 %) haemoglobin solution extracted from horse blood with the extinction coefficient of human oxygenated haemoglobin shows strong similarity in the visible and NIR range (Fig. 4A), as does deoxygenated haemoglobin (Fig. 4B, sO_2_ = 0 %). The range of measured *µ_a_* exceeds the upper limit of detection for the Shimadzu SIS, resulting in detector saturation truncating the Soret band (406 nm).^56^ Additionally, noise artefacts from silicon to InGaAs detector changeover betweeen 850 – 900 nm have been removed by our data processing. The oxygenated haemoglobin solution yields a very low SAM score of 0.03198 ± 0.00002 rad relative to our H_2_O characterisation when compared with Prahl,^52^ while the spectral angle of the deoxygenated haemoglobin solution is relatively high at 0.25670 ± 0.00108 rad (Table 2). This is likely due to the use of sodium dithionite for deoxygenation, which can form methaemoglobin that has characteristic absorption peaks at 404, 508 and notably 635 nm,^57^ as present in our spectrum.

**Fig 4.**
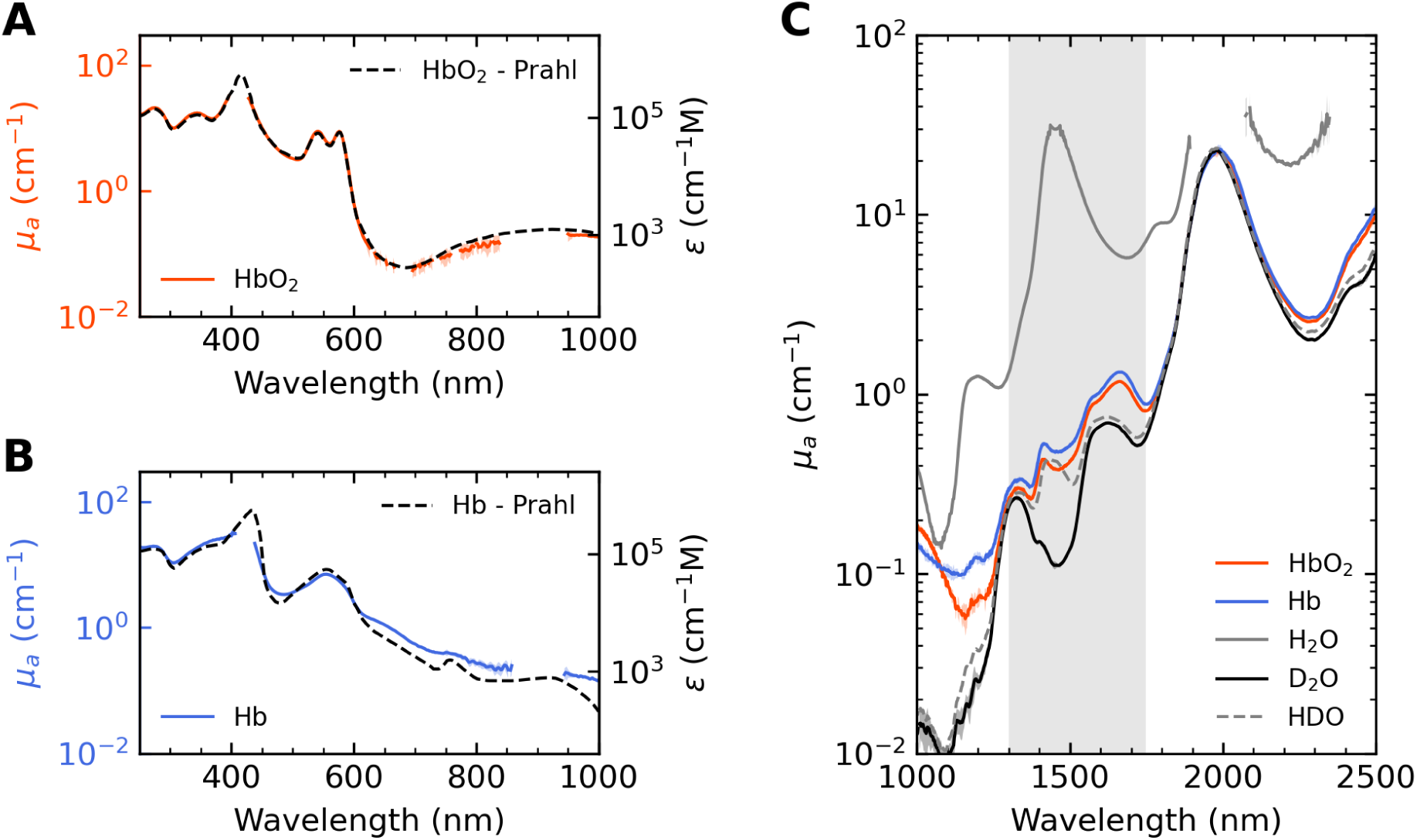
Characterisation of the absorption coefficient *µ_a_* of oxygenated and deoxygenated haemoglobin extracted from whole horse blood from 250 – 2500nm. A) Estimated *µ_a_* of oxygenated horse haemoglobin in D_2_O (solid red) compared with the extinction coefficient of oxygenated human haemoglobin compiled by Prahl^52^ (dashed black). B) Estimated *µ_a_* of deoxygenated haemoglobin in D_2_O (solid blue) compared with the extinction coefficient of deoxygenated human haemoglobin compiled by Prahl^52^ (dashed black). C) Short-wave infrared characterisation of the haemoglobin solutions in (A) and (B) compared with H_2_O (solid grey), D_2_O (solid black) and a 1% solution of H_2_O in D_2_O (HDO, dashed grey).

SWIR characterisation of haemoglobin (Fig 4C) shows absorption peaks between 1300 – 1750 nm, however, these overlap with H_2_O and D_2_O features. HDO, semiheavy water, is a naturally occurring molecule containing one hydrogen, one deuterium and one oxygen atom^58^ – the peaks observed suggest the presence of HDO in our samples, which could have arisen from exposure of our D_2_O to air over time or some H_2_O remaining in our D_2_O-washed haemoglobin pellet. Features between 1300 – 1750 nm have also been observed in prior work,^28^ and do not appear to be haemoglobin-specific absorption characteristics due to consistency with HDO absorption features.

Additionally, the isosbestic point at 1100 nm is likely not physiological, but an artefact of sample processing that could be attributed to the presence of methaemoglobin increasing NIR absorption.^57^

### 3.4 Optical properties of lard and corn oil agree with literature

Estimated *µ_a_* of corn oil and lard (Fig. 5A) were compared with a reference for pig fat by van Veen *et al.*^50^ between 429 – 1098 nm and a reference for human fat by Anderson *et al.*^24^ covering 1123 – 2319 nm. The lard and corn oil spectra are both highly consistent with these references, producing SAM scores of 0.11875 ± 0.01109 rad for lard and 0.08582 ± 0.00152 rad for corn oil between 1000 – 1600 nm (Table 2). As lipid absorption approaches the lower limit of detection of our sensors in the visible and NIR, *µ_a_* estimation failed between 500 – 900 nm. Consequently, statistical analysis was not performed in the visible and NIR.

**Fig 5.**
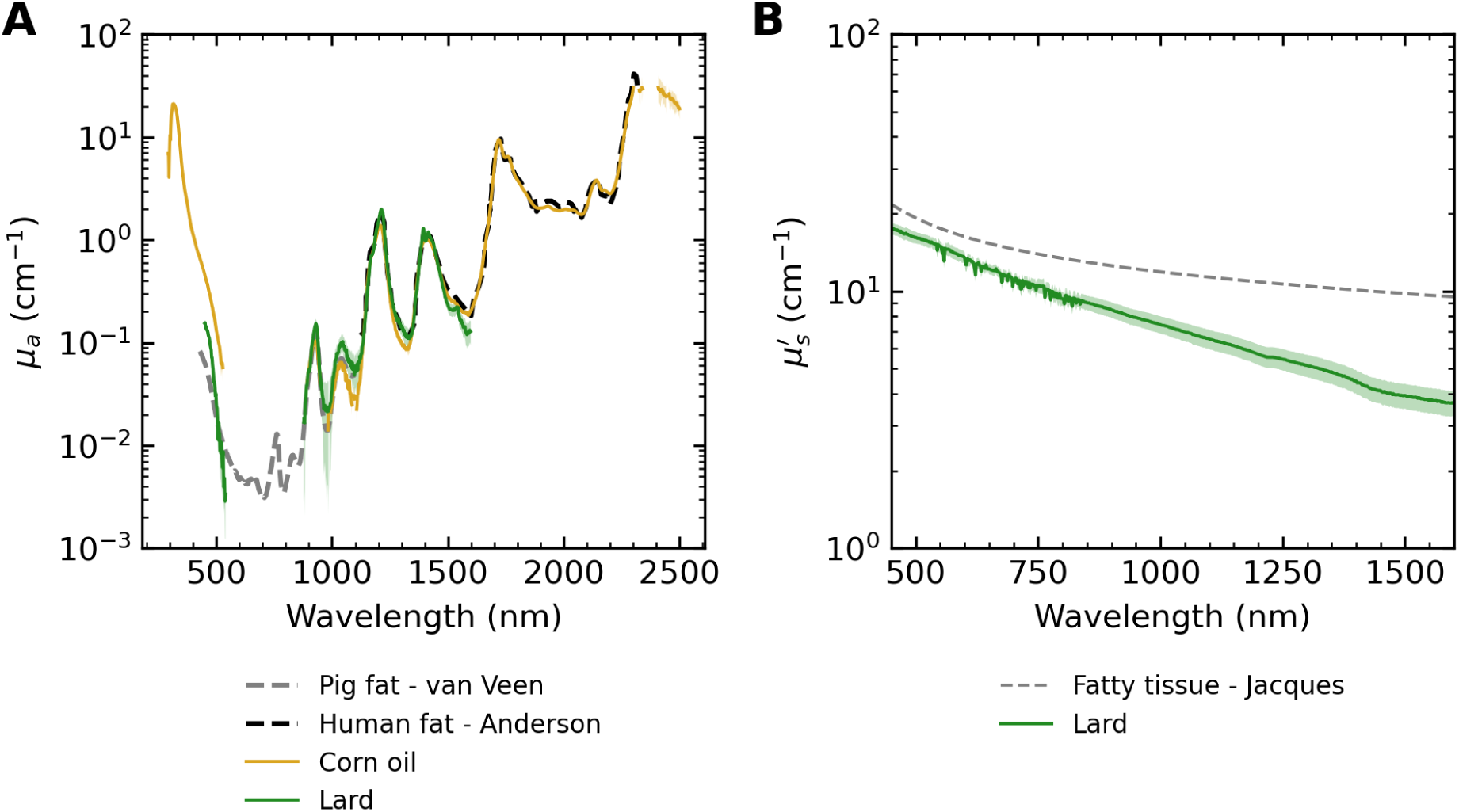
Characterisation of the absorption and reduced scattering coefficients (*µ_a_* and *µ_s_*′, respectively) of lard from 450 – 1600 nm and *µ_a_* of corn oil from 300 – 2500 nm. A) *µ_a_* of solid lard (reduced pig fat, solid green) and corn oil (solid gold) compared with pig fat measured by van Veen *et al.*^50^ (dashed grey) and human fat by Anderson *et al.*^24^ (dashed black). B) *µ_s_*′ of solid lard (solid green) compared with a model of fatty tissue generated using Rayleigh and Mie theory using coefficients compiled by Jacques^36^ extrapolated to 1600 nm (dashed grey).

We compared our *µ_s_*′ characterisation of lard with a model of *µ_s_*′ for fatty tissue generated using Rayleigh and Mie theory^36^ extrapolated to 1600 nm (Fig. 5B). Our estimated *µ_s_*′ of lard shows an inverse power law decay, consistent with Mie theory scattering but distinct from the fatty tissue model. Whilst our lard is lower in scattering amplitude than the fatty tissue model, differences are small at visible and NIR wavelengths, suggesting that the Jacques model might not be valid for extrapolation to SWIR. The observed differences could also be attributed to lard being a reduced pig fat (99.8 % fat), thus having different particle size and concentration to *in vivo* fatty tissue.

### 3.5 Synthetic melanin shows a steep decline in SWIR absorption

Our *µ_a_* characterisation of synthetic melanin dissolved in DMSO (Fig. 6A) shows an inverse power law decay until 1100 nm, beyond which absorption by DMSO dominates the spectrum, suggesting low melanin absorption at SWIR wavelengths. To further investigate synthetic melanin absorption at SWIR wavelengths, we subtracted DMSO absorption from the synthetic melanin in DMSO spectrum (solid brown) and scaled the result to match the melanosome absorption spectrum approximation by Jacques^36^ (dashed black) at 500 nm (Fig. 6B). The melanosome absorption approximation is an exponential function fitted to melanosome interior measurements from laser experiments^59^ (black stars). Here, strong divergence between the melanosome approximation, which we have extrapolated into the SWIR, and our measured synthetic melanin absorption is observed in the NIR through to the 1400 nm water absorption peak. Quantitatively, the scaled, DMSO subtracted synthetic melanin spectrum compared to Jacques’ melanosome approximation yields a strong spectral angle of 0.08451 ± 0.00721 rad between 450 – 1000 nm, but strong discordance between 1000 – 1600 nm with a high spectral angle of 0.35036 ± 0.02198 rad as well as a low *R* of 0.715 (confidence interval = [0.643, 0.774]) (Table 2). Absorption peaks characteristic of the synthetic melanin solution occur around 1200 nm, 1500 nm and 1900 nm (shaded grey bars, Fig. 6A and B), likely due to OH group vibrational overtones^60^ as the observed features coincide with H_2_O absorption peaks (dashed grey, Fig. 6B).

**Fig 6.**
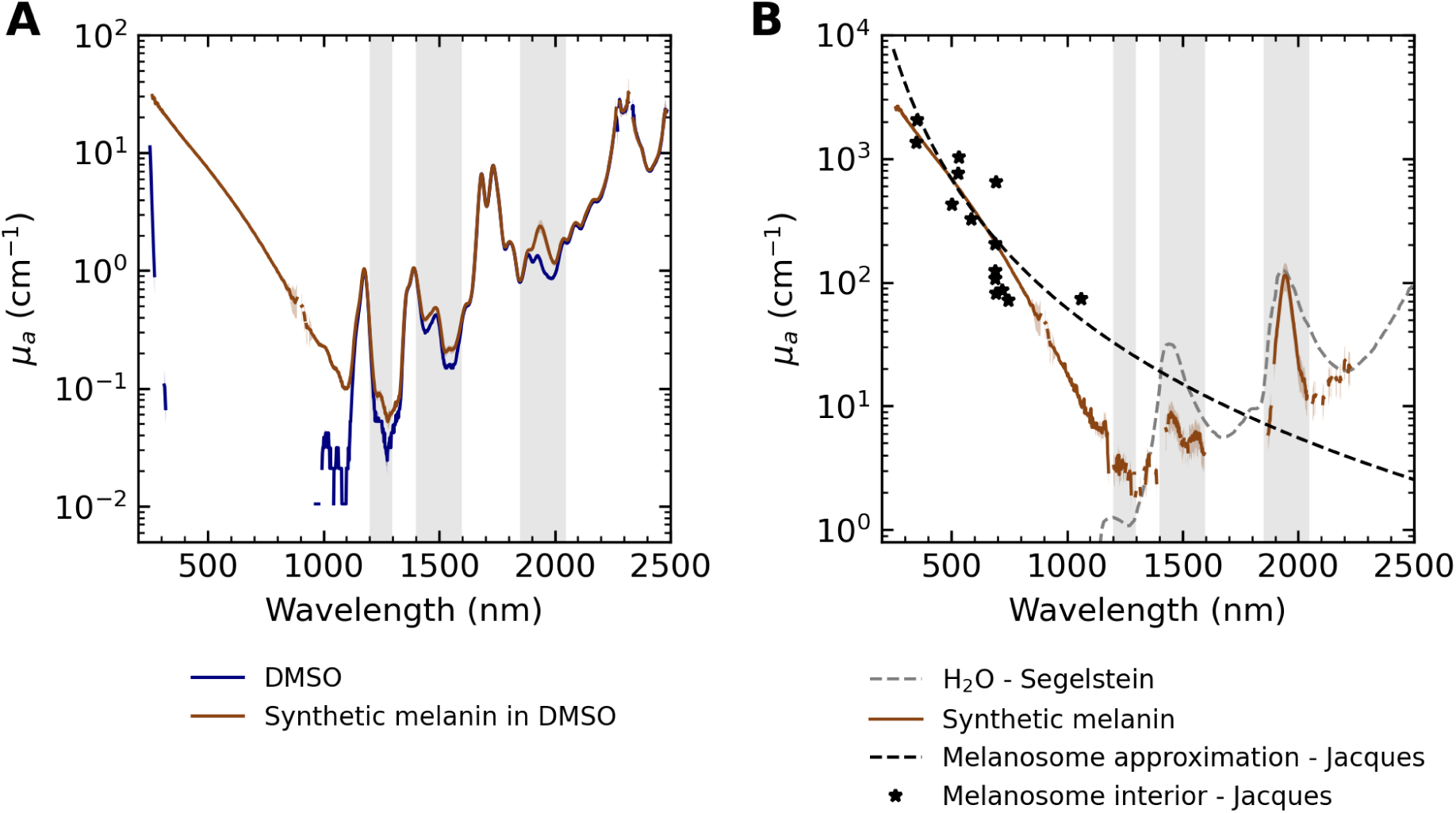
Characterisation of the absorption coefficient (*µ_a_*) of synthetic melanin. A) Estimated *µ_a_* of synthetic melanin dissolved in dimethyl sulfoxide (DMSO, 0.21 mg/mL, solid brown) and DMSO (solid navy). Absorption is below the limit of detection for our detector for the pure DMSO spectrum between 300 – 1000 nm. B) DMSO subtracted synthetic melanin *µ_a_* spectrum (solid brown) scaled to match melanosome absorption spectrum approximation by Jacques^36^ (dashed black) at 500 nm, derived from measurements of the melanosome interior by Jacques^59^ (black stars). We note absorption peaks in the synthetic melanin solution around 1200 nm, 1500 nm and 1900 nm (shaded grey bars in (A) and (B)) that we attribute to OH group vibrational overtone modes.^60^

### 3.6 Comparison of simulated tissues with tissue mimicking phantoms

A dataset for continuous visible to SWIR tissue simulations comprised of spectra collected in this work (solid lines) and spectra sourced from SIMPA’s libraries (dashed lines, scattering spectra extrapolated to 1600 nm) is presented (Fig. 7A and B). Models of fat and muscle built from this dataset were probed using Monte Carlo simulations and compared with the fabricated tissue mimicking phantoms (Fig. 7E).

**Fig 7.**
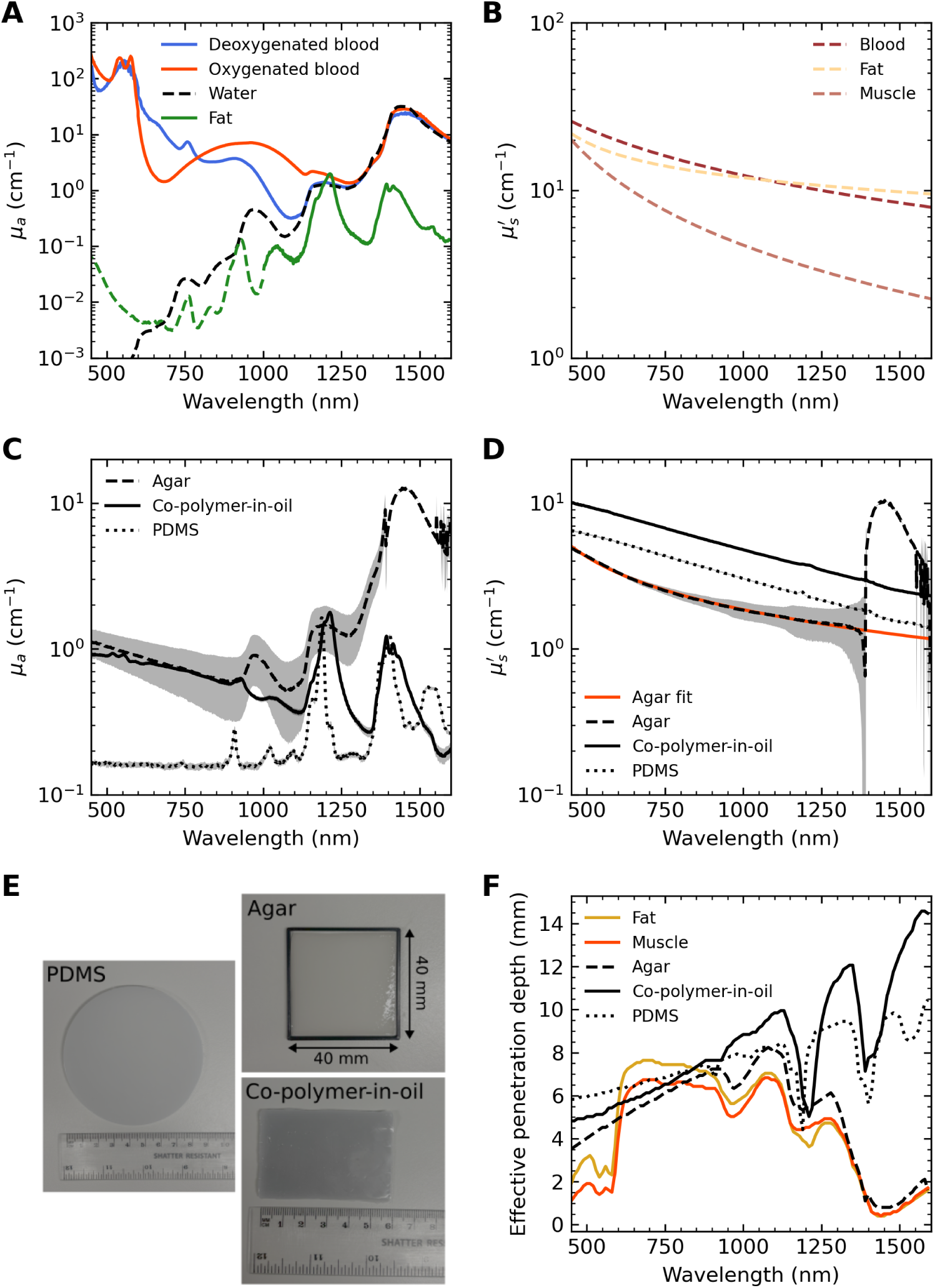
Evaluating the suitability of tissue mimicking materials in the SWIR. A) Dataset of absorption coefficients (*µ_a_*) required for modelling muscle and fat. Measured *µ_a_* of oxygenated and deoxygenated whole blood (solid red and blue, respectively) and lard (solid green) are collated with *µ_a_* references for pig fat^50^ (dashed green) and water^23^ (dashed black). B) Dataset of reduced scattering coefficients (*µ_s_*′) for soft tissues modelled using Rayleigh and Mie theory using coefficients collated by Jacques^36^ and Gröhl,^29^ extrapolated to 1600 nm. C and D) Measured *µ_a_* (C) and *µ_s_*′ (D) spectra of agar (dashed black), PDMS (dotted black) and copolymer-in-oil (solid black) phantoms. E) Exemplary photographs of the fabricated agar, PDMS and co-polymer-in-oil phantoms. F) Effective penetration depth of the fat (solid yellow) and muscle (solid orange) tissue simulations compared with simulations of the agar (dashed black), PDMS (dotted black) and co-polymer-in-oil (solid black) phantoms.

Tissue phantom optical properties show consistent absorption in the visible and NIR by the agar (dashed black) and copolymer-in-oil (solid black) phantoms, which diverges in the SWIR due to the water content of the agar dominating SWIR absorption (Fig. 7C). In contrast, the PDMS material (dotted black) shows low absorption in the visible and NIR with SWIR absorption comparable with the copolymer-in-oil (Fig. 7C). In terms of scattering (Fig. 7D), the PDMS (dotted black) and co-polymer-in-oil (solid black) phantoms show an inverse power law relationship, while the agar (dashed black) presents an exponential decay with a strong peak at 1400 nm, consistent with the water absorption peak as shown in Fig. 7C. This feature is thus assumed to be an artefact and a fit to Eq. (8) of the agar *µ_s_*′ spectrum cropped to 1350 nm was calculated (solid red) for use in the agar Monte Carlo model (Fig. 7D).

Monte Carlo simulations of the tissue phantoms, constructed from the measured phantom optical properties (Fig. 7C and D), were compared with our fat and muscle models using effective penetration depth, calculated as the depth at which fluence drops to 1*/e* of its maximum value (Fig. 7E). Aside from the oxygenated haemoglobin peaks between 500 – 600 nm, the fat (solid yellow) and muscle (solid orange) simulations show similar light propagation to the agar, copolymer-in-oil and PDMS (dashed, solid and dotted black, respectively) in the visible and NIR. In SWIR, the agar phantom is most consistent with the fat and muscle simulations due to their high water contents, resulting in shallow light penetration between 1350 – 1600 nm where water absorption is high. In contrast, the co-polymer-in-oil and PDMS phantoms have much deeper SWIR penetration.

## 4 Discussion

The short-wave infrared wavelength range holds promise for opening new avenues in biomedical optics, however, more data characterising the optical properties of tissue chromophores are needed to assess the capabilities of these wavelengths. Here, we characterised the optical properties of the dominant chromophores in tissue (water, haemoglobin, melanin and lipids) and commonly-used tissue-mimicking materials using single-integrating-sphere systems. We reported *µ_a_* and *µ_s_*′ for whole blood and lard, and *µ_a_* only for the non-scattering chromophores: H_2_O, D_2_O, oxygenated and deoxygenated haemoglobin, corn oil and synthetic melanin. The resulting spectra were integrated with an open-source Python toolkit^29^ for continuous visible to SWIR optical tissue simulations. Computational models of muscle and fat were probed with Monte Carlo simulations and validated against tissue mimicking phantoms.

Our characterisation of whole horse blood revealed a comparable magnitude of oxygenated whole blood absorption between 700 – 1000 nm and 1000 – 1300 nm, suggesting that haemoglobin absorption could be targeted by SWIR optical sensors. We note low absorption by deoxygenated blood between 1000 – 1300 nm, yielding a higher contrast between oxygenated and deoxygenated blood absorption in this range than previously reported.^41^ To decouple contributions from H_2_O, we measured solutions of haemoglobin in D_2_O, where similarly to our whole blood measurements, haemoglobin absorption was distinguishable from the D_2_O solvent between 1000 – 1300 nm. Beyond 1300 nm, haemoglobin absorption was indistinguishable from H_2_O, HDO and D_2_O absorption features for the whole blood and haemoglobin solution samples. The processing required to prepare isolated oxygenated and deoxygenated haemoglobin samples reconstituted purely in an alternative solvent is challenging. Contrary to our whole blood characterisations, our haemoglobin in D_2_O characterisations were highly sensitive to sample processing, resulting in the presence of HDO and an isosbestic point at 1100 nm. Isosbestic points near 1200 nm have been observed in prior work by Gruensfelder *et al.*^28^ (from whom we based our protocol from), highlighting the delicate nature of these characerisations and the need for more studies analysing the absorption properties of haemoglobin in the SWIR.

Our characterisation of synthetic melanin showed lower absorption in the SWIR than the conventional extrapolation of the melanosome absorption approximation for skin by Jacques.^36^ The melanosome approximation equation is an exponential curve fitted to measurements of the melanosome interior (biological melanin) in laser experiments^59^ of which the highest wavelength is 1064 nm, thus, measurements at longer wavelengths are required to validate extrapolation of this curve to SWIR. Our findings show an inverse power law decay, which is valid initially, but begins to diverge from Jacques’ approximation in the NIR. Lower melanin absorption in the SWIR could be harnessed in biomedical optics to mitigate the confounding effects of skin tone on biomedical data.^25^ The correspondence of this lower melanin absorption with strong contrast observed between oxygenated and deoxygenated blood around 1100 nm suggests promise for the implementation of SWIR wavelengths in oximetry. We used synthetic melanin, a brown pigment produced by the oxidation of tyrosine with hydrogen peroxide, as an analogue for biologically produced melanin. Further studies of biological melanins in melanosomes, rather than analysing freely-diffusing synthetic melanin, would be valuable for interpreting our observations in the context of the scattering contributions introduced by melanosome structure and the absorption properties of biological melanins.

Our measurements of the absorption coefficient of corn oil and lard compared favourably with references for fat by van Veen^50^ and Anderson *et al.*^24^ between 1000 – 1600 nm. We manually extracted Anderson’s data points using plot digitising software^61^ since these absorption spectra were not tabulated. We note that Anderson presents the *µ_a_* of water and fat on the same figure, with different *x* axes. We have observed studies^62^ which report these lipid data with a blue shift of around 80 nm in the SWIR, aligned with the water *x* axis. Since Anderson’s measurements are one of the few SWIR lipid spectra available in literature, this example highlights the need for open, tabulated datasets to ensure reproducibility.

In optical simulations, *µ_s_*′ is often modelled using Rayleigh and Mie theory, fitting around measurements of *µ_s_*′ at 500 nm.^36^ These models are quoted to be valid for 400 – 1300 nm,^36^ however, in this work we extrapolated out to 1600 nm to compare with our SWIR measurements and run SWIR tissue simulations. The need for more data at longer wavelengths to identify an accurate model for scattering at longer wavelengths has been highlighted in literature.^25, 36^ We observed divergence of our reduced scattering spectrum for lard with a modelled fatty tissue spectrum at longer wavelengths, suggesting that empirical observations would be valuable to inform more accurate models of scattering in tissues. The observed difference could be explained by differences in tissue structure or particle size of lard (reduced fat) and *in vivo* fatty tissue, or stronger contributions from Mie theory at longer wavelengths as we approach the sample particle size. Further studies measuring the scattering properties of tissues at SWIR wavelengths should be conducted to validate these models.

We constructed models of muscle and fat to demonstrate how our dataset of measured optical properties can be implemented to explore SWIR light-tissue applications. To evaluate the suitability of tissue-mimicking phantoms for SWIR applications, we fabricated, characterised and simulated three highly cited recipes^46–48^ for commonly used phantom materials, tuned with dyes and scatterers to mimic soft tissue: agar (with added India ink and Intralipid), PDMS (with added India ink and titanium dioxide) and co-polymer-in-oil (with added nigrosin and titanium dioxide). We selected effective penetration depth as a metric to compare photon propagation through the fat, muscle and phantom models probed with Monte Carlo simulations. Our study showed broad agreement of the muscle and fat models with the tissue-mimicking phantoms in the visible and NIR. In SWIR, the agar phantom closely matched the penetration depth of the muscle and fat simulations due to the high water content of these materials. In contrast, significantly deeper photon penetration was achieved with the PDMS and co-polymer-in-oil simulations beyond 1200 nm, indicating that these materials (which, unlike agar, are stable and reusable) would benefit from dyes mimicking the strong absorption of the 1400 nm water absorption peak to be suitable SWIR tissue proxies. Copolymer-in-oil, however, remains a good mimic for highly fatty tissues below the 1400 nm water absorption peak as the mineral oil base provides the 1200 nm absorption peak.

## 5 Conclusion

The optical absorption and reduced scattering coefficients of the dominant tissue chromophores: water, haemoglobin, lipids, and melanin, were characterised from visible to SWIR wavelengths. These data were integrated into an open-source Python toolkit to create a dataset of optical properties for SWIR tissue simulations. The optical properties of tissue-mimicking phantoms, including agar, PDMS and copolymer-in-oil based materials, were measured and compared with simulated soft tissues, revealing the limitations of current phantom recipes in mimicking SWIR light-tissue interactions. Overall, open and consistent datasets of optical properties will enable robust and reproducible tissue simulations, facilitating rapid assessment and prototyping of next-generation spectroscopy and imaging techniques.

## Code and Data Availability

All data and code used in the preparation of this paper and not otherwise referenced inline will be made available on the University of Cambridge Apollo repository (https://doi.org/10.17863/CAM.132008) upon paper acceptance or Github (https://github.com/BohndiekLab/SWIR-tissue-chromophore-characterisations), respectively.

## Acknowledgments

This work was supported by the Engineering and Physical Sciences Research Council Centre for Doctoral Training in Sensor Technologies for a Healthy and Sustainable Future [EP/S023046/1].

We would like to acknowledge funding support from Cancer Research UK under grant number C9545/A29580 (SEB, TRE), the UKRI Engineering and Physical Sciences Research Council (EPSRC) under grant numbers EP/X037770/1 (SEB, LM) and APP16931 (SEB, IR). This project has received funding from the European Research Council (ERC) under the European Union’s Horizon 2020 research and innovation programme (project NEURAL SPICING grant agreement No. 101002198 (LM, TRE). RT acknowledges the financial support of the Trinity Barlow Scholarship, Cambridge Trust International Scholarship and the Herchel Smith Scholarship. JG acknowledges funding from the Walter Benjamin Stipendium of the Deutsche Forschungsgemeinschaft. This work was supported, in whole or in part, by the Gates Foundation [INV-077759] (MJW, LM, SEB). The conclusions and opinions expressed in this work are those of the author(s) alone and shall not be attributed to the Foundation. Under the grant conditions of the Foundation, a Creative Commons Attribution 4.0 License has already been assigned to the Author Accepted Manuscript version that might arise from this submission. Please note works submitted as a preprint have not undergone a peer review process. We thank the Cancer Research UK Cambridge Institute Flow Core and Research Instrumentation and Cell Services Core for their support in conducting this research. We would also like to thank Lorna Wright, Aparajita Naik, Steve Mead and Darren Robyler for their support in conducting this research.

## Biographies

**Melissa J. Watt** is pursuing her PhD in physics at the University of Cambridge. She received her MPhys degree in physics from the University of Bath in 2022 and her MRes degree in Sensor Technologies for a Healthy and Sustainable Future from the University of Cambridge in 2023. She is researching quantitative optical spectroscopy techniques for applications in equitable, non-invasive optical monitoring and diagnostics.

**Ran Tao** completed her MRes in connected electronic and photonic systems in 2021 and is currently pursuing her PhD in physics, both from the University of Cambridge. Her research focuses on broadband systems for estimating tissue optical absorption and scattering properties, extending optical measurements from near-infrared-I (650 to 950 nm) to short-wavelength infrared (1000 to 2500 nm).

**Sarah E. Bohndiek** completed her PhD in radiation physics at the University College London in 2008 and then worked in both the United Kingdom (at Cambridge) and the United States (at Stanford) as a postdoctoral fellow in molecular imaging. Since 2013, she has been a group leader at the University of Cambridge, where she is jointly appointed in the Department of Physics and the Cancer Research UK Cambridge Institute. She was appointed as a full professor of biomedical physics in 2020. Sarah was elected Fellow of the international optics and photonics society SPIE in 2020.

Biographies and photographs of the other authors are not available.

## List of Figures

1. **Absorption and scattering coefficients (*µ****_a_* **and *µ****_s_***) of dominant tissue chromophores can be estimated from visible to short-wave infrared wavelengths from measurements of transmittance and reflectance using single integrating sphere (SIS) systems.** A) Dominant chromophores in tissue. Created in BioRender. Bohndiek, S. (2026) https://BioRender.com/urkzunj B) Schematic of a single integrating sphere and cuvettes showing transmittance and reflectance. Adapted from Tao *et al.*^38^
2. **Estimations of the absorption coefficient** *µ_a_* **of H_2_O and D_2_O are consistent with references from literature from 1000 – 1600 nm.** Estimated *µ_a_* of H_2_O (A) and D_2_O (B) calculated using Beer-Lambert analysis of transmittance measurements from the Shimadzu SIS (solid red) are compared to reference data (dashed grey) by Segelstein^23^ (A) and Kedenburg^34^ (B). Estimated *µ_a_* of H_2_O measured by Tao, calculated with reflectance and transmittance measurements from the in-house SIS using the inverse adding doubling algorithm, are presented (dashed black).
3. **Characterisation of the absorption coefficient (***µ_a_***) of oxygenated and deoxygenated whole horse blood from 450 – 1600 nm and reduced scattering coefficient (***µ_s_*′**) from 600 – 1600 nm.** A) Estimated *µ_a_* of a 1% solution of whole blood in phosphate-buffered saline (PBS) is presented from 450 – 600 nm, stitched to estimated *µ_a_* of undiluted whole blood from 600 – 1600nm (solid red). Similarly, a 1% solution of deoxygenated whole blood in PBS is presented from 450 – 600nm, stitched to the estimated *µ_a_* of undiluted whole blood from 600 – 1600nm (solid blue). An average *µ_a_* of oxygenated (dashed red) and deoxygenated (dashed blue) human whole blood compiled from literature references by Bosschaart *et al.*^41^ is compared with our estimations for *µ_a_*. B) Estimated *µ_s_*′ of oxygenated and deoxygenated whole horse blood from 600 – 1600nm (solid red and blue, respectively). An average *µ_s_*′ compiled from literature by Bosschaart *et al.*^41^ (dashed black) and an approximation based on Rayleigh and Mie theory using coefficients optimised by Gröhl *et al.*^29^ (dashed grey). C) Brightfield images of horse red blood cells (RBCs) imaged with flow cytometer. Four population types were identified side profile RBCs (A), circular cells with a concave middle (B, assumed to be erythrocytes), flat cells (C, assumed to be spherocytes) and star-shaped cells (D, assumed to be Burr cells). The cyan overlays show the masking function used to measure cell diameter by the flow cytometer software. D) Histogram showing distribution of horse RBC diameters from all RBC populations (black) with population B overlayed (green).
4. **Characterisation of the absorption coefficient *µ_a_*** of oxygenated and deoxygenated haemoglobin extracted from whole horse blood from 250 – 2500nm. A) Estimated *µ_a_* of oxygenated horse haemoglobin in D_2_O (solid red) compared with the extinction coefficient of oxygenated human haemoglobin compiled by Prahl^52^ (dashed black). B) Estimated *µ_a_* of deoxygenated haemoglobin in D_2_O (solid blue) compared with the extinction coefficient of deoxygenated human haemoglobin compiled by Prahl^52^ (dashed black). C) Short-wave infrared characterisation of the haemoglobin solutions in (A) and (B) compared with H_2_O (solid grey), D_2_O (solid black) and a 1% solution of H_2_O in D_2_O (HDO, dashed grey)
5. **Characterisation of the absorption and reduced scattering coefficients (***µ_a_* **and** *µ_s_*′ **, respectively) of lard from 450 – 1600 nm and** *µ_a_* **of corn oil from 300 – 2500 nm.** A) *µ_a_* of solid lard (reduced pig fat, solid green) and corn oil (solid gold) compared with pig fat measured by van Veen *et al.*^50^ (dashed grey) and human fat by Anderson *et al.*^24^ (dashed black). B) *µ_s_*′ of solid lard (solid green) compared with a model of fatty tissue generated using Rayleigh and Mie theory using coefficients compiled by Jacques^36^ extrapolated to 1600 nm (dashed grey).
6. **Characterisation of the absorption coefficient (***µ_a_***) of synthetic melanin.** A) Estimated *µ_a_* of synthetic melanin dissolved in dimethyl sulfoxide (DMSO, 0.21 mg/mL, solid brown) and DMSO (solid navy). Absorption is below the limit of detection for our detector for the pure DMSO spectrum between 300 – 1000 nm. B) DMSO subtracted synthetic melanin *µ_a_* spectrum (solid brown) scaled to match melanosome absorption spectrum approximation by Jacques^36^ (dashed black) at 500 nm, derived from measurements of the melanosome interior by Jacques^59^ (black stars). We note absorption peaks in the synthetic melanin solution around 1200 nm, 1500 nm and 1900 nm (shaded grey bars in (A) and (B)) that we attribute to OH group vibrational overtone modes.^60^
7. **Evaluating the suitability of tissue mimicking materials in the SWIR.** A) Dataset of absorption coefficients (*µ_a_*) required for modelling muscle and fat. Measured *µ_a_* of oxygenated and deoxygenated whole blood (solid red and blue, respectively) and lard (solid green) are collated with *µ_a_* references for pig fat^50^ (dashed green) and water^23^ (dashed black). B) Dataset of reduced scattering coefficients (*µ_s_*′) for soft tissues modelled using Rayleigh and Mie theory using coefficients collated by Jacques^36^ and Gröhl,^29^ extrapolated to 1600 nm. C and D) Measured *µ_a_* (C) and *µ_s_*′ (D) spectra of agar (dashed black), PDMS (dotted black) and co-polymer-in-oil (solid black) phantoms. E) Exemplary photographs of the fabricated agar, PDMS and co-polymer-in-oil phantoms. F) Effective penetration depth of the fat (solid yellow) and muscle (solid orange) tissue simulations compared with simulations of the agar (dashed black), PDMS (dotted black) and co-polymer-in-oil (solid black) phantoms.

## List of Tables

1. Coefficients for modelling reduced scattering coefficient of tissues using Eq. (8). ^∗^*a*^′^ for blood and muscle converted from *µ_s_* to *µ_s_*′ using Eq. (1) assuming *g* = 0.98 and *g* = 0.9 for blood and muscle, respectively.
2. Spectral angle mapping score (SAM), Pearson correlation coefficient (*R*), and root mean square error (RMSE) of measured absorption coefficients compared against references from literature.

